# *HOXA9* acts as a regulatory switch in acute myeloid leukaemia and myeloproliferative neoplasms

**DOI:** 10.1101/2021.03.05.434116

**Authors:** Laure Talarmain, Matthew A. Clarke, David Shorthouse, Jasmin Fisher, Benjamin A Hall

**Author notes:** The authors declare no competing financial interests.

## Abstract

Blood malignancies arise from the dysregulation of haematopoiesis. The type of blood cell and the specific order of oncogenic events initiating abnormal growth ultimately determine the cancer subtype and subsequent clinical outcome. HOXA9 plays an important role in acute myeloid leukaemia (AML) prognosis by promoting blood cell expansion and altering differentiation; however, the function of HOXA9 in other blood malignancies is still unclear. Here, we demonstrate the importance of this gene in chronic myeloproliferative neoplasms (MPN) and highlight the biological switch and prognosis marker properties of HOXA9 in AML and MPN. This binary switch function can explain the clinical stratification of these two blood disorders. First, we establish the ability of HOXA9 to stratify AML patients with distinct cellular and clinical outcomes. Then, through the use of a computational network model of MPN, we show that the self-activation of HOXA9 and its relationship to JAK2 and TET2 can explain the branching progression of JAK2/TET2 mutant MPN patients towards divergent clinical characteristics. Finally, we predict a connection between the RUNX1 and MYB genes and a suppressive role for the NOTCH pathway in MPN diseases.

## Introduction

Blood cancers are malignancies that can arise from any type of blood cell and dramatically affect haematopoiesis. Myeloproliferative Neoplasms (MPNs) are chronic diseases of the myeloid lineage characterised by an excessive production of fully functional terminally differentiated blood cells. These have been classified into 3 types: polycythemia vera (PV), essential thrombocythemia (ET), and primary myelofibrosis (PMF) [1]. Despite the relatively good prognosis of these diseases, MPN patients are at high risk of thrombosis and can develop a blast phase MPN (MPN-BP) [2]; a subtype of the blood cancer Acute Myeloid Leukemia (AML) with poor survival outcomes [3]. The frequency of MPN transformation to blast phase MPN is highly related to the initial MPN disease type [4,5,6]. Therefore, a better understanding of the molecular events driving the different subtypes of MPNs is essential to help diagnose patients with higher risk of thrombosis and AML progression.

AML itself is an aggressive blood and bone marrow malignancy defined by the uncontrolled growth of myeloid progenitor cells along with a myeloid-lineage differentiation arrest [7]. As with MPN, there exist different types of AML with a broad range of morphologic, cytogenic and immunologic features, all associated with diverse clinical outcomes [8]. Despite their similarities; prognosis, symptoms, and genetic alterations differ between AML and MPN. For example, *JAK2* mutation is the main driver event of MPN diseases yet is rarely found in *de novo* AML [9]. However, myeloid lineage dysregulation by both MPN and AML, as well as the ability of MPN to evolve to AML, indicate that both diseases may share similar biological mechanisms. The identification of these processes will help identify aberrant genes and pathways in blood malignancies that could be targets for new drugs.

Better understanding of the patterns of genetic alterations in cancer cells can also be used for the classification of blood diseases and prediction of progression into more severe forms of the disease [10]. How different combinations and orders of mutations lead to different subtypes of cancer remains a major open question [11, 12]. The importance of mutation order has been demonstrated in MPN by Ortmann et al [13], who show that two subpopulations of patients with MPN can be distinguished by the order of mutation acquisition between the *TET2* and *JAK2* genes and that these subpopulations have distinct clinical characteristics. Further analyses of these cohorts show that patients with *JAK2* mutated before *TET2* are younger at presentation of the disease in clinics, are more likely to present with PV, have a higher risk of thrombosis and respond better to JAK2 inhibitor ruxolitinib. However, the molecular interplay between both mutations within cancer cells and how their order rather than their combination triggers dissimilar clinical characteristics have not been investigated.

Overexpression of a single homeobox gene, *HOXA9*, has been reported as sufficient to quickly induce myeloproliferation, gradually followed by AML progression after a period of time [14]. Homeobox genes or HOX genes were first identified in the fruit fly *Drosophila melanogaster* as essential regulators of early embryogenesis [15] and are thought to have a critical role in cancer development [16]. In the HOXA family, *HOXA9* is the most described gene in literature and its expression was shown to be the single most highly correlating factor, out of 6817 genes tested, for poor prognosis in AML [17]. The importance of *HOXA9* in AML has been widely explored; however, this has mainly focused on specific AML subtypes such as MLL-rearranged leukaemia [18] and NUP98-*HOXA9* induced leukaemia [19], while its role in other blood malignancies such as MPN or other AML subtypes is poorly characterised. Recently, the oncogenic property of *HOXA9* has been associated with its self-positive feedback loop in myeloid precursor cells as a result of its ability to bind its own promoter [20]. We hypothesise in this work that this specific property can help stratify patients with blood cancers affecting the myeloid lineage.

Using public datasets from AML patients and MPN studies, we show that bimodal *HOXA9* expression identifies two distinct cohorts of patients/mice, reflecting the protein acting as a binary switch in the cell. We show that this switch-like behaviour enables a clinical stratification in these blood diseases leading to separation of individuals into two clinically distinct populations. This leads to distinct prognoses, but also allows specific disease type classification. First, *HOXA9* bimodal expression affects the clinical features, such as age and white blood cell counts, but also patient classification into specific French-American-British (FAB) or molecular subtypes. Next, we design a computational network model that offers a mechanistic explanation of the distinct clinical features of MPN progression in patients with different orders of *JAK2* and *TET2* mutations. This computational model predicts that *HOXA9* is directly downstream of *JAK2* and *TET2* and effectively stores their mutational history, leading to a phenotypic switch in double mutant cells dependent upon mutation order and producing distinct subtypes of the disease. Finally, the network model also predicts a suppressive role for the NOTCH pathway in MPN and a new interaction between *RUNX1* and *MYB*.

## Materials and Methods

### Analysis and visualisation of public cancer datasets

The AML patient data was generated by TCGA and downloaded with Firebrowse (RNAseq). The file contains data from 173 patients. We use the Transcripts Per Million (TPM) normalised data (TCGA RNAseq) and the R package multimode [21] to determine the significance of gene expression bimodality. The *modetest* command is used to reject unimodality with the default ACR method and a p-value at 0.05. We use the R package Survival [22] and the AML clinical data from TCGA to plot the survival curves and compute the p-values of the log rank test. Overall survival variables are in months and left as is in the analyses. We use the Plotly Python Open Source Graphing library to plot Sankey diagrams (available at https://plot.ly).

### Differentially expressed genes in HOXA9 cohorts

We use a python script to separate patients between the two *HOXA9* expression peaks found in the AML data from TCGA. 31 patients constitute the low peak and 80 the high peak. We compute the absolute difference of the mean expression of each gene between each cohort to find the genes which are most differentially expressed between the two groups of patients. We rank the genes from the highest to the lowest differential expression and take the first 30 genes from this list.

This workflow was repeated in the comparison of the two *HOXA9* cohorts. The same arbitrary cut-off, 30 first genes with the highest fold change, mostly identified in the HOX family or genes with no determined role in haematopoiesis.

### Microarray data analysis

The microarray data set reported in [23] is available in the ArrayExpress repository at European Molecular Biology Laboratory–European Bioinformatics Institute (http://www.ebi.ac.uk/arrayexpress/) and is accessible through the ArrayExpress accession number E-MTAB-2986. While 12 samples are described in the paper, only 11 could be found in the public data, with one wild type sample missing in the microarray.

For analysis, from the set of all transcripts in the microarray, the genes with a low detection p-value (below 0.05) were filtered and transformed with a quantile normalisation. The ComplexHeatmap R library was used to plot the heatmaps [24].

### XGBoost

XGBoost (e**X**treme **G**radient **Boost**ing) was used to rank different gene pathways that have been well described in cancer to identify which pathways and genes amongst these pathways have the highest correlation with a gene of interest and its expression level in the AML patients (TCGA RNASeq) [25]. More details can be found in the Supplementary Information.

### Executable network model of MPN

Computational models of MPN cancer fate determination were constructed as a qualitative network (QN) in the BioModelAnalyzer [26]. This process is described in more detail in Supplemental methods, but briefly QNs are constructed from reported gene interactions in the wider literature, and refined by testing model behaviour against reported phenotypes.

## Results

### 1- *HOXA9* expression separates cohorts of AML patients with distinct clinical features

Ectopic expression of *HOXA9* in AML has been widely demonstrated, but few studies have investigated the biological attributes of this transcription factor contributing to leukemogenesis. Zhong et al [20] have shown that *HOXA9* in cell lines can induce its own expression through a positive feedback loop, which promotes a continuous differentiation block and self-renewal leading to increase of hematopoietic stem cells and development of leukaemia. To validate *HOXA9* self-activation and the oncogenic role in leukaemia in patients, we studied its expression in AML patient data from The Cancer Genome Atlas (TCGA) [27]. We find that *HOXA9* has bimodal expression in these data (Fig. 1A). This bimodality separates patients into two cohorts: 31 patients in the low expression peak, and 80 patients in the high expression peak. A survival analyses of both groups using Kaplan-Meier survival curves and the log-rank test confirms that *HOXA9* can be used as a marker of poor prognosis in AML (Fig. 1B). This patient stratification based upon *HOXA9* expression is consistent with the reported positive feedback loop characteristic of this gene and suggests that once activated or inhibited, the gene would maintain its expression level, leading to divergence in the disease progression.

**Figure 1.**
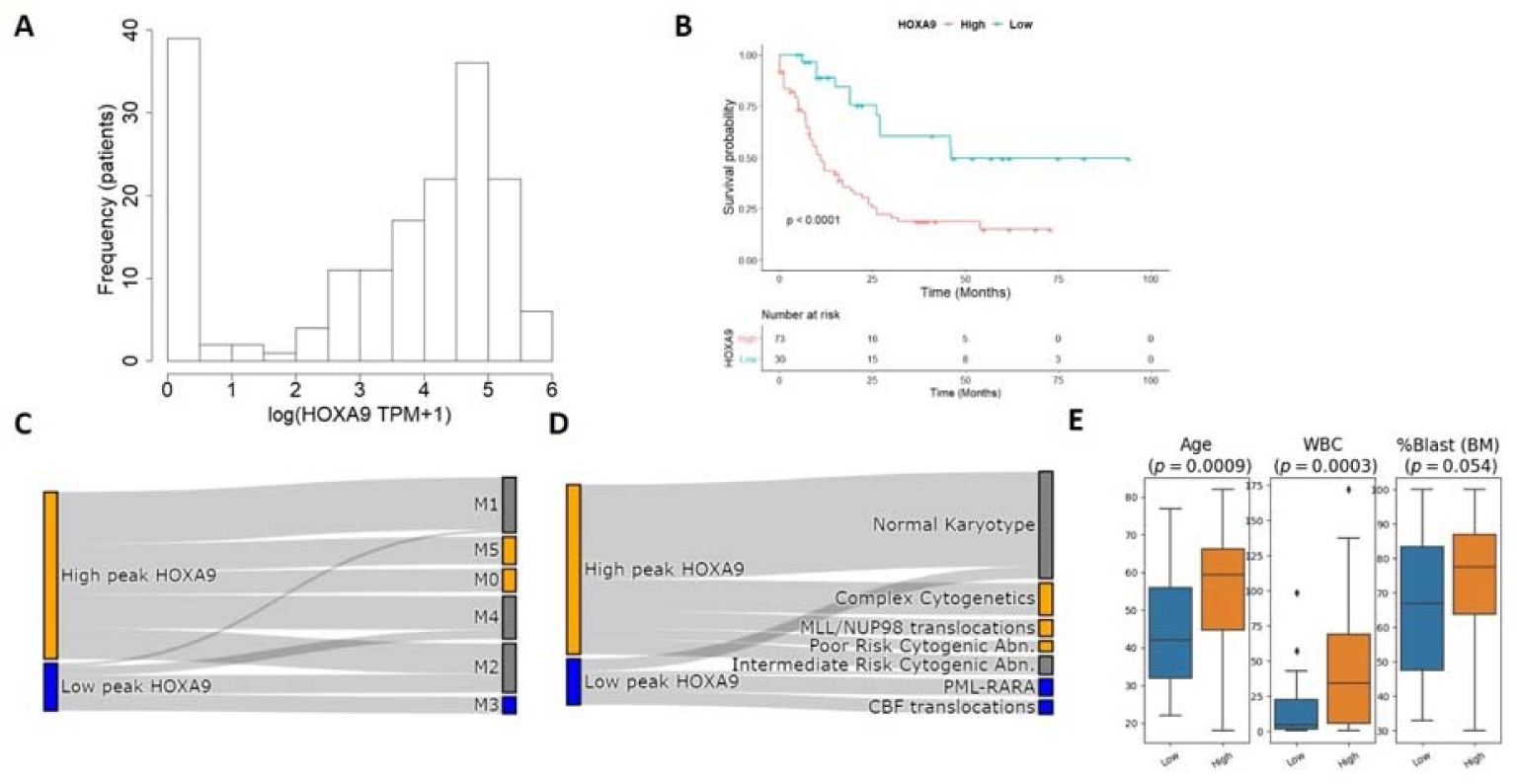
*HOXA9* low and high expression stratifies patients in AML. **(A)** *HOXA9* expression in AML patients is significantly bimodal (unimodality test rejected with p < 2.2 × 10^−16^), suggesting a role as a genetic switch. **(B)** *HOXA9* high and low cohorts have divergent AML prognosis, consistent with known *HOXA9* biology in AML. *HOXA9* expression also partially explains **(C)** FAB and **(D)** molecular classifications of AML patients. Using Sankey diagrams, we find that the M3 FAB subtype as well as the PML-RARα and CBF (CBFB-MYH11 and RUNX1-RUNXT1) translocations are solely linked to low expression of *HOXA9*. Similarly, the high cohort possesses specific AML subclasses: M0, M5, MLL/NUP98 translocations and complex cytogenetics. **(E)** *HOXA9* cohorts show distinct clinical characteristics: high cohort patients are older, display a higher white blood cell counts (WBC) and tend to have higher percentage of blasts in the bone marrow.

To investigate if the switch role property of *HOXA9* impacts AML subtypes, we looked at the distribution of French-American-British (FAB, named M0-M7), and molecular classifications among the two *HOXA9* cohorts. We show that different *HOXA9* expression cohorts exclude specific FAB subtypes (Fig. 1C). This suggests that *HOXA9* expression is strongly coupled to FAB subtypes.

In light of these findings, we looked to characterise the common features of *HOXA9* expression cohorts. Cytogenic aberrations and gene rearrangements are frequent in AML and are known to alter the disease morphology as well as the clinical features and prognosis [27]. We find that *HOXA9* stratifies patients with different molecular classification (Fig. 1D). MLL-induced leukaemia has been linked to high *HOXA9* [18], while M3 AML subtype is characterised by PML-RARα translocation and low *HOXA9* in the literature [28]. Low *HOXA9* expression in AML with RUNX1-RUNXT1 and CBFB-MYH11 abnormalities, which constitute the core binding factor (CBF) AML, was also established but unexplained in literature [29]. With this work, we confirm these findings and further establish the correlation between high *HOXA9* expression and the M0 and M5 subtypes as well as complex cytogenetics. Finally, we expand the *HOXA9* stratification effect by looking at other available clinical features in the data and find that *HOXA9* expression correlates with age, white blood cell count (WBC) and blast percentage in the bone marrow (Fig. 1E).

PML-RARα, RUNX1-RUNXT1 (AML1-ETO) or CBFB-MYH11 chromosomal abnormalities confer good prognosis in AML patients [30, 31]. All these aberrations are linked to low *HOXA9* expression which also exhibits good survival prognosis among patients compared to high expression. To confirm that high *HOXA9* is a poor prognosis marker independently of its associated molecular aberrations or FAB subtypes, we look at survival outcomes within FAB classes. M0, M3 and M5 being all specific to one cohort, we examine the survival of patients within the M2 and M4 subtypes for high and low *HOXA9* expression. Survival curves and log rank tests within both subtypes confirm the poor prognosis marker function of high *HOXA9* (Fig. S1 and S2). These results as well as the diverging clinical features among patient cohorts are consistent with *HOXA9* becoming trapped in high- or low- expression states through self-activation in AML diseases.

### 2- The *JAK2*/*TET2*/*HOXA9* motif explains divergent disease clinical outcomes in MPN

*JAK2* is the most commonly mutated gene in many MPN patients, but different subtypes of the disease with distinct clinical traits are observed [32]. In contrast, *TET2* was only recently identified in blood studies. First discovered in MPN in 2008 by Delhommeau et al [33], *TET2* mutation resulting in its function loss has been associated with diverse hematologic malignancies [34]. We have shown that *HOXA9* can enable clinical stratification in AML through its positive feedback loop.

Ortmann [13] describes a bifurcation among MPN patients that acquire *JAK2* and *TET2* mutations in different orders. This raises the question whether *HOXA9* expression could also explain this divergent clinical symptoms.

To address this question, we constructed a computational network model in a multistep process. In order to reproduce the branching in MPN patients, the underlying network of gene interactions must include genes that are sensitive to the mutation order [35]. This requires that parts of the network act a switch, capable of storing “memory” of previous events. This “memory” property can be encoded by a positive feedback loop acting on a gene that is downstream of both mutated genes [36]. This hypothetical gene must additionally respond differently to each of the mutations. That is to say, one mutation must activate the gene whilst the other inhibits it, so that the gene can maintain its change in activity after the occurrence of the second mutation. The loop is necessary to induce this inheritable change in the presence of constitutive reset processes such as protein and RNA degradation.

We developed a computational model of this simple gene motif with *JAK2* and *TET2* genes and a hypothetical gene target with a positive feedback loop (Fig. 2A). We identified HOXA9 as a gene with connections to *TET2* and *JAK2* that was consistent with our computational motif. *TET2* and *JAK2* have been indirectly and directly linked to *HOXA9* activity (Fig. 2B). *STAT5* is a well-known downstream target of *JAK2* [37], and it is also established that *STAT5* and *HOXA9* act as binding partners in hematopoietic cells [38]. Furthermore, it was recently shown that tyrosine phosphorylation of *HOXA9* is *JAK2*-dependent [39] and seems to increase the effect of *HOXA9* on its downstream targets [39]. Regarding the interaction of *TET2* with *HOXA9*, Bocker et al found significantly reduced expression of *HOXA* genes when *TET2* expression is lost [40]. In particular *HOXA9* expression in kidney is significantly decreased by *TET2* loss. *HOXA9* is therefore activated by both *JAK2* and *TET2* and possesses a self-positive feedback loop property [20]. Therefore, the *JAK2*/*TET2*/*HOXA9* motif shares all the required properties for observing a clinical divergence in blood diseases.

**Figure 2.**
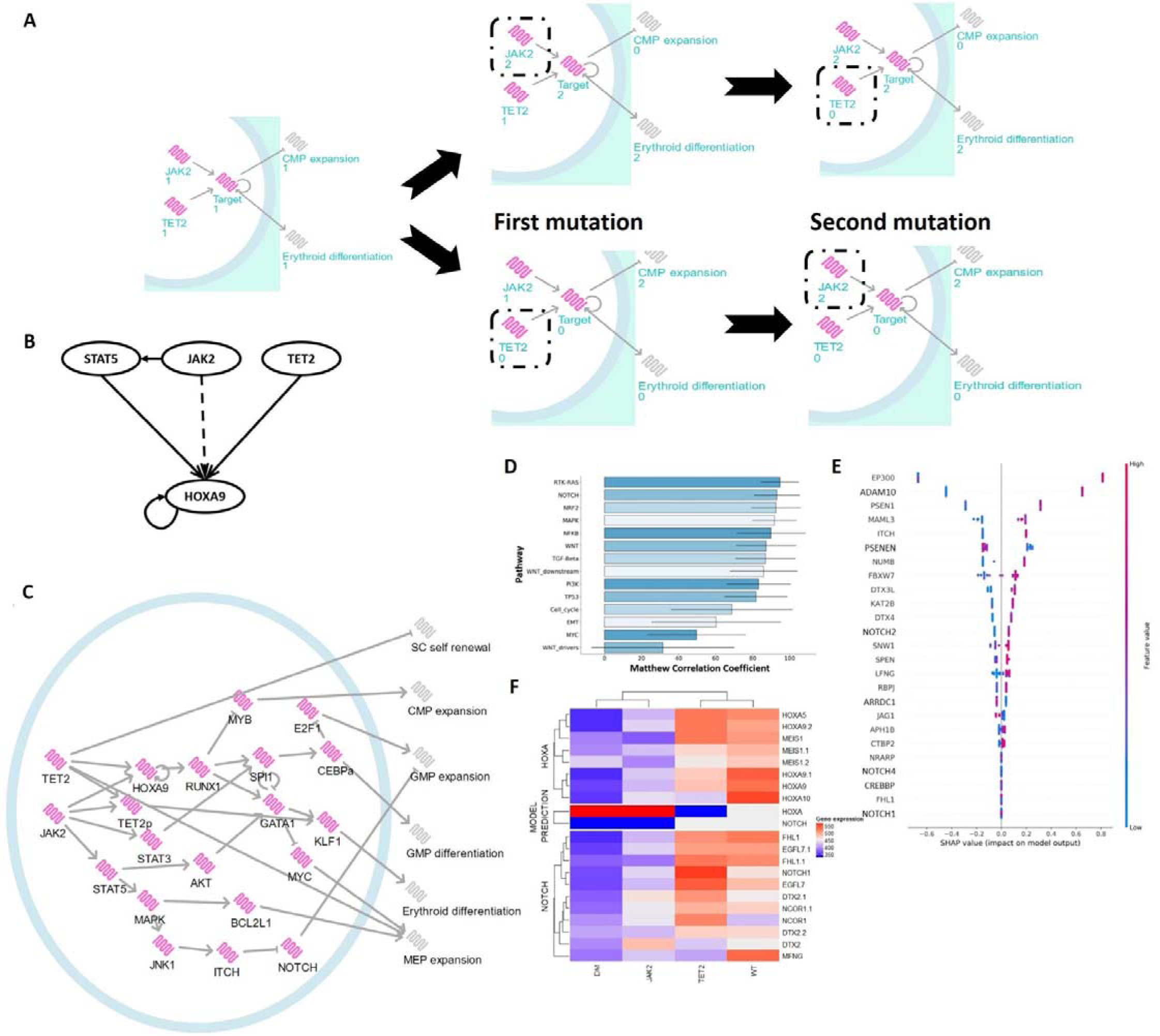
*HOXA9* determines cancer cell fates in MPN diseases with *JAK2* and *TET2* mutations. **(A)** MPN patients with *JAK2*/*TET2* mutations showing different clinical characteristics can be explained by a simple gene motif including a switching property. The model starts from a healthy state on the left (wild type) and sequentially acquires mutations in *JAK2* and *TET2* genes. The first mutation affects the gene target expression (middle networks) which remains stable when the second mutation appears. The order in which mutations occur impacts on the gene target expression but also the phenotypes, CMP expansion and erythroid differentiation (networks on the right). Despite the identical final mutational state, the cell shows different hematopoietic behaviours. **(B)** *HOXA9* is downstream of both *TET2* and *JAK2* and acts as a developmental switch through its self-positive feedback loop. Direct interaction drawn as continuous arrows and post translation modification as dotted arrows. **(C)** The *JAK2*/*TET2/HOXA9* molecular network is built with the BioModelAnalyzer (BMA) tool and integrates six phenotypes considered as our model outputs. Using the Linear Temportal Logic model checking tool available in BMA, we observe a bifurcation after simulation of the double mutant state. The bifurcation analysis identifies two stable states with different phenotype values that fit the mutation order characteristics observed in MPN patients with *JAK2* and *TET2* mutations. **(D)** Ranking of major cancer pathways in *JAK2* analyses determine the RTK-RAS pathway as the most correlated to *JAK2*. This pathway contains *JAK2* and so this result is as expected. The second pathway is NOTCH. The overall accuracy of the pathway is computed with the Matthews Correlation Coefficient. **(E)** We plot SHAP scores for the NOTCH pathway to determine which genes of the pathway have been important classifiers, that is genes with an important expression correlation with *JAK2*. Among them, *PSEN1* and *ADAM10* are involved in the cleavage of NOTCH membrane receptors. *NUMB* activates *ITCH* which degrades NOTCH. **(F)** The heatmap of the NOTCH pathway and HOXA family generated using MPN microarray datasets from [23] validate NOTCH expression in our model highlighting the importance of this pathway in MPN disease progression. HOXA heatmap confirms HOXA bimodality but show different levels of expression to what we find in our model. “JAK2” and “TET2” refer to the single mutant mouse models, and “DM” is the double mutant with *JAK2* mutated first. “WT” designates the wild type (no mutation) genotype.

Based on this *JAK2*/*TET2*/*HOXA9* motif, we refine our computational model to reproduce the observed biological differences between patients with different combinations of *JAK2* and *TET2* mutations. To do so, we extend our computational network model featuring three main genes: *JAK2*, *TET2*, and *HOXA9* (Fig. 2C). Our model includes six phenotypes relevant to cancer development: stem cell self-renewal, common myeloid progenitor (CMP) expansion, granulocyte-monocyte progenitor (GMP) expansion, GMP differentiation, erythroid differentiation and megakaryocyte-erythroid progenitor (MEP) expansion. To capture the wider biology of MPN progression, we further included important hematopoietic markers in our computational model. A detailed literature review and full description of how we built the network model are available in Table S1, Table S2 and the Supplemental Methods.

Finally, four fundamental cancer genotypes are defined: the wild type (no mutation), the *TET2* single mutant, the *JAK2* single mutant and the double mutant (in either order) (Table 1). The wild type model illustrates haematopoiesis in its healthy state. The single mutants are defined using the literature (see Supplementary Information). The final genotype is the double mutant which can lead to one of two cancer endpoints (fixpoint attractors that represent one of the two clinical outcomes). Each fixpoint represents either *TET2* first or *JAK2* first double mutants and are defined from results presented by Ortmann et al [13]. Our computational model as shown in Figure 2C reproduces the expected behaviours described in Table 1 and therefore the clinical stratification observed in Ortmann et al [13]. The model suggests that the elevated differentiation observed in the *JAK2* first double mutants [13] is induced by the increased expression of *RUNX1*, *KLF1* and *GATA1* as well as the downregulation of *MYC* not found in *TET2* first double mutant. This gene expression difference between double mutants can partly explain the divergent clinical behaviours between the two groups of patients, including the increased risk of thrombosis and the faster diagnosis as a result of the abnormally high number of differentiated cells in these patients.

**Table 1.** Specification table for *JAK2*/*TET2* BMA model. In order from the left to the right columns, the specifications are the wild type state, the *TET2* single mutant, the *JAK2* single mutant and finally the double mutants, which consists of a bifurcation with two state attractors that represent the case where *TET2* is mutated before *JAK2* (*TET2* first) and the alternative case where *JAK2* is mutated first. We determine phenotype values using literature for the single mutants and Ortmann et al [13] for the double mutants. The value 1 represents the healthy state, 0 the lowered/inactive state and 2 the overactive state.

|  | WT | *TET2* | *JAK2* | *TET2*<br>first | *JAK2*<br>first |
| --- | --- | --- | --- | --- | --- |
| Stem Cell<br>Renewal | 1 | 2 | 1 | 2 | 2 |
| CMP expansion | 1 | 2 | 1 | 2 | 1 |
| GMP expansion | 1 | 2 | 2 | 2 | 2 |
| GMP<br>differentiation | 1 | 0 | 1 | 1 | 1 |
| Erythroid<br>differentiation | 1 | 0 | 2 | 1 | 2 |
| MEP expansion | 1 | 1 | 2 | 2 | 2 |

### 3- MPN network predicts new gene dynamics and interactions

The computational model identifies new gene dynamics as part of the MPN disease progression. In the complete network model, *HOXA9* requires both *JAK2* and *TET2* expression to remain active (Table S2). Upregulation of either *JAK2* or *HOXA9* results in the hyperactivation of *HOXA9* while *TET2* loss causes inactivation. Wild type activity is maintained by the balance of these two genes. JAK2 activation mutation and TET2 loss both drive the system into a committed state. JAK2 activation raises HOXA9 activity to a level at which it can maintain its activity through control of its own expression. Subsequent loss of TET2 does not impact its activity as this hyperactivation makes it independent of TET2. Conversely, TET2 loss causes a loss of HOXA9 expression in the cell, rendering it insensitive to subsequent JAK2 activation. This occurs as HOXA9 expression drops, preventing a subsequent response to JAK2 activation due to low concentration of the protein in the cell. Therefore, a possible explanation of the order dependence could be a combination of mutual dependency between *JAK2* and *TET2* to activate *HOXA9*, combined with the positive feedback self-loop of *HOXA9* itself.

Whilst building single mutant phenotypes, we noticed a relationship between *JAK2* and GMP expansion is required to match the increased number of myeloid progenitors observed in organisms with a *JAK2* mutation. To explore possible pathways downstream of *JAK2* that could explain this link to myeloid diseases, we applied a machine learning approach (XGBoost) to AML TCGA data as a relevant and closely related blood cancer. We found that *JAK2* is highly correlated with the NOTCH pathway (Fig. 2D), which has been found to act as a tumour suppressor in leukemia due to the great expansion of GMP cells after loss of NOTCH signalling [41]. From the SHAP scores of NOTCH genes plotted in Figure 2E, we identified *ITCH* to be among the genes with a high score in the NOTCH pathway for *JAK2*. *ITCH* controls the degradation of NOTCH [42] and is found to be induced by JNK1 [43] from the MAPK pathway which is a well-known downstream pathway of *JAK2*/*STAT5* [44]. We therefore suggest that the *JAK2* path to GMP expansion could be MAPK and NOTCH dependent.

Another new interaction predicted by our network model is the inhibition of *MYB by RUNX1*. The Common Myeloid Progenitors (CMP) are found to be differentially expanded between *JAK2* and *TET2* first patients in Ortmann et al [13]. In our initial model, we integrated an inhibition interaction between *SPI1* and *MYB*, our CMP expansion marker, a connection which has been found experimentally [45]. This inhibition and the stable *SPI1* expression in the double mutant states prevented the known bifurcation in CMP expansion in double mutants. Further investigations lead us to suggest that the bifurcation could be obtained by replacing *SPI1* by *RUNX1* for the *MYB* inhibition which is supported by different studies. *RUNX1* activates *SPI1* and *GATA1*, and both are found to be inhibitors of *MYB* [45, 46]. Additionally, conditional knockout of *RUNX1* in mice results in enhanced CMP frequencies [47][48]. All together, these findings suggest that *RUNX1* can be linked to CMP expansion via *MYB* inhibition.

### 4- Analyses of public MPN datasets validate NOTCH role in MPN as well as HOXA9 bimodality and prognosis role in blood diseases

To validate the predictions arising from our MPN computational model, we compared our findings to public MPN data not used in the model construction. Chen et al compare MPN with different *JAK2* and *TET2* mutational profiles using transcriptomic mouse data [23]. We compare the gene expression of pathways/gene subsets to those we have included in our model to determine if our model fits their data. We first find that the NOTCH pathway behaves as predicted (Fig. 2F). We also find that the trend in the expression of *RUNX1*, *MYC* and *MYB* support our model (Fig. S3, S4 and S5). Confusingly *HOXA9* expression showed a “switching” behaviour in this mouse model but associated a low expression of *HOXA9* with *JAK2* mutations and high expression with *TET2* mutations.

Jeong et al [49] previously demonstrated the direct phosphorylation of *TET2* by *JAK2* in a combination of *in vitro* human/murine hematopoietic cell lines with erythroid characteristics. Once phosphorylated *TET2* activates *KLF1*, an important positive regulator of erythropoiesis [50]. In this context, loss of *TET2* implies reduced erythroid differentiation which is in agreement with our model. In the same study, the authors show in a murine cell line that *JAK2* mutation leads to *HOXA9* upregulation. These findings are consistent with our *JAK2*/*TET2*/*HOXA9* motif but disagree with Chen’s microarray experiments where *HOXA9* expression is lowered in *JAK2* single and double mutants (Fig. 2F). Given the downstream genes follow the expected expression, this raises the question of whether the interactions in the original motif should be replaced by a pair of inhibitions expected by Chen data rather than activations. Whilst there exist possible routes to connect *TET2* and *HOXA9* through an inhibition, and to change the sign of the downstream interactions for *HOXA9*, we are however unable to find evidence of inhibition of *HOXA9* by *JAK2*. We further note that as the Jeong data is human derived, it may be a more representative experimental model system. Both datasets however support the role of *HOXA9* as a binary switch in MPN.

## Discussion

Out of 6817 genes tested *HOXA9* is the single most highly correlating factor for poor prognosis due to treatment failure in AML [17]. This finding has made *HOXA9* the most studied gene in the HOXA family. Here we have demonstrated that it acts in AML as a discrete switch rather than a spectrum, which impact AML clinical characteristics such as classification and survival. Our study shows that despite the existing correlation between *HOXA9* expression and some AML subtypes, its poor prognosis characteristic is independent of patient classification. We further extend the prognosis marker role of the *HOXA9* gene to another blood disorder, MPN. We show that in MPN diseases with *JAK2*/*TET2* mutations, *HOXA9* high expression is found in the *JAK2* first patients while *TET2* first patients display lower *HOXA9* expression. As *JAK2* first patients have a higher risk of developing thrombosis compared to *TET2* first patients, and as thrombotic events are the main causes of death in MPNs patients [51], this further suggests a deleterious influence of *HOXA9* high expression on patient clinical outcomes in another myeloid disease and emphasise the role of *HOXA9* as a poor prognosis marker in blood malignancies.

In addition to providing new insights into the regulatory control of cancer cell fate through *HOXA9*, our computational network model recapitulates the disease symptoms using well-known hematopoietic transcription factors. Further investigations of these genes could benefit clinicians by designing new drugs or applying already existing treatments to reduce symptoms and the risk of developing blast phase MPN. In addition to the specific claims of the model, several other clinical implications arise. One key feature of *TET2*-first MPN patients is their reduced sensitivity to Ruxolitinib, a JAK2 inhibitor drug [13]. Interestingly our computational model suggests that after *TET2* loss, most common *JAK2* targets are unchanged by *JAK2* activation mutation due to the “switching” property exerted by *HOXA9* self-loop. It follows that *JAK2* inhibition is therefore inefficient for those genes. Finally, whilst *JAK2* is the main driver mutation found in all MPN patients, different diseases with distinct clinical traits can be observed [32]. Until now, the source of this clinical diversity following *JAK2* mutation was unclear. Here, we demonstrate that patients who first had a *TET2* mutation have a reduced number of erythroid cells as a result of *TET2* indirect downregulation of *GATA1* and *KLF1* which explain the reduced number of PV diseases in *TET2* first patients despite the presence of *JAK2* mutation [13]. While *JAK2* dysregulation may be the principal driver of MPNs, other mutations shape the disease clinical type by altering the normal development of distinct hematopoietic subpopulations. Finally, we predict the involvement of the NOTCH pathway in MPN diseases. NOTCH shows both oncogenic and tumour suppressor roles in different tissues and in the hematopoietic system: NOTCH favours cancer growth in T acute lymphoblastic leukemia (T-ALL) through its *MYC* activation but is also found to augment the host immune response against cancer by activation of M1 macrophages [52]. The role of NOTCH in hematopoietic stem and progenitor cells is still an on-going debate, however, it seems that a certain level of NOTCH signalling is required to protect individuals from hematological malignancies [53]. We suggest that *JAK2* increases GMP expansion through its inhibitory effect on NOTCH via the MAPK pathway and ITCH and so predict a tumour suppressor role for NOTCH in the GMP cell population. Our network model suggests a novel mechanism for understanding how cancer fate can be determined through regulatory switches and highlights several new areas for further studies.

## Acknowledgements

We thank members of the Hall group at the MRC-Cancer Unit and the Fisher group at the University College London Cancer Institute, and Aleksandra Watson at the University of Cambridge for valuable discussions. BAH acknowledges support from the Royal Society (grant no. UF130039). DS and BAH were further supported by the Medical Research Council (grant no. MR/S000216/1) LT acknowledges support from Microsoft Research. JF was supported by the National Institute for Health Research University College London Hospitals Biomedical Research Centre and Cancer Research UK.

## Competing Interests

The authors declare no competing financial interests.

## Notes

### Competing Interest Statement

The authors have declared no competing interest.

## References

[1] J. L. Spivak, “Myeloproliferative neoplasms,” New England Journal of Medicine, vol. 376, no. 22, pp. 2168–2181, 2017.

[2] A. Tefferi, M. Mudireddy, F. Mannelli, K. H. Begna, M. M. Patnaik, C. A. Hanson, R. P. Ketterling, N. Gangat, M. Yogarajah, V. De Stefano et al., “Blast phase myeloproliferative neoplasm: Mayo-agimm study of 410 patients from two separate cohorts,” Leukemia, vol. 32, no. 5, pp. 1200–1210, 2018.

[3] M. Yogarajah and A. Tefferi, “Leukemic transformation in myeloproliferative neoplasms: a literature review on risk, characteristics, and outcome,” in Mayo Clinic Proceedings, vol. 92, no. 7. Elsevier, 2017, pp. 1118–1128.

[4] A. Tefferi, P. Guglielmelli, D. R. Larson, C. Finke, E. A. Wassie, L. Pieri, N. Gangat, R. Fjerza, A. A. Belachew, T. L. Lasho et al., “Long-term survival and blast transformation in molecularly annotated essential thrombocythemia, polycythemia vera, and myelofibrosis,” Blood, The Journal of the American Society of Hematology, vol. 124, no. 16, pp. 2507–2513, 2014.

[5] A. Tefferi, E. Rumi, G. Finazzi, H. Gisslinger, A. Vannucchi, F. Rodeghiero, M. Randi, R. Vaidya, M. Cazzola, A. Rambaldi et al., “Survival and prognosis among 1545 patients with contemporary polycythemia vera: an international study,” Leukemia, vol. 27, no. 9, pp. 1874–1881, 2013.

[6] T. Barbui, J. Thiele, F. Passamonti, E. Rumi, E. Boveri, M. Ruggeri, F. Rodeghiero, E. S. d’Amore, M. L. Randi, I. Bertozzi et al., “Survival and disease progression in essential thrombocythemia are significantly influenced by accurate morphologic diagnosis: an international study,” Journal of clinical oncology, vol. 29, no. 23, pp. 3179–3184, 2011.

[7] C. S. Grove and G. S. Vassiliou, “Acute myeloid leukaemia: a paradigm for the clonal evolution of cancer?” Disease models & mechanisms, vol. 7, no. 8, pp. 941–951, 2014.

[8] U. Bacher, W. Kern, S. Schnittger, W. Hiddemann, C. Schoch, and T. Haferlach, “Further correlations of morphology according to fab and who classification to cytogenetics in de novo acute myeloid leukemia: a study on 2,235 patients,” Annals of hematology, vol. 84, no. 12, pp. 785–791, 2005.

[9] J. Aynardi, R. Manur, P. R. Hess, S. Chekol, J. J. Morrissette, D. Babushok, E. Hexner, H. J. Rogers, E. D. Hsi, E. Margolskee et al., “Jak2 v617f-positive acute myeloid leukaemia (aml): a comparison between de novo aml and secondary aml transformed from an underlying myeloproliferative neoplasm. a study from the bone marrow pathology group,” British journal of haematology, vol. 182, no. 1, pp. 78–85, 2018.

[10] R. C. Lindsley, “Uncoding the genetic heterogeneity of myelodysplastic syndrome,” Hematology 2014, the American Society of Hematology Education Program Book, vol. 2017, no. 1, pp. 447–452, 2017.

[11] D. G. Kent and A. R. Green, “Order matters: the order of somatic mutations influences cancer evolution,” Cold Spring Harbor perspectives in medicine, vol. 7, no. 4, p. a027060, 2017.

[12] M. A. Clarke, S. Woodhouse, N. Piterman, B. A. Hall, and J. Fisher, “Using state space exploration to determine how gene regulatory networks constrain mutation order in cancer evolution,” in Automated reasoning for systems biology and medicine. Springer, 2019, pp. 133–153.

[13] C. A. Ortmann, D. G. Kent, J. Nangalia, Y. Silber, D. C. Wedge, J. Grinfeld, E. J. Baxter, C. E. Massie, E. Papaemmanuil, S. Menon et al., “Effect of mutation order on myeloproliferative neoplasms,” New England Journal of Medicine, vol. 372, no. 7, pp. 601–612, 2015.

[14] C. Bach, S. Buhl, D. Mueller, M.-P. Garcá-Cuéllar, E. Maethner, and R. K. Slany, “Leukemogenic transformation by hoxa cluster genes,” Blood, The Journal of the American Society of Hematology, vol. 115, no. 14, pp. 2910–2918, 2010.

[15] E. B. Lewis, “A gene complex controlling segmentation in drosophila,” in Genes, development and cancer. Springer, 1978, pp. 205–217.

[16] S. Bhatlekar, J. Z. Fields, and B. M. Boman, “Hox genes and their role in the development of human cancers,” Journal of molecular medicine, vol. 92, no. 8, pp. 811–823, 2014.

[17] T. R. Golub, D. K. Slonim, P. Tamayo, C. Huard, M. Gaasenbeek, J. P. Mesirov, H. Coller, M. L. Loh, J. R. Downing, M. A. Caligiuri et al., “Molecular classification of cancer: class discovery and class prediction by gene expression monitoring,” science, vol. 286, no. 5439, pp. 531–537, 1999.

[18] J. Faber, A. V. Krivtsov, M. C. Stubbs, R. Wright, T. N. Davis, M. van den Heuvel-Eibrink, C. M. Zwaan, A. L. Kung, and S. A. Armstrong, “Hoxa9 is required for survival in human mll-rearranged acute leukemias,” Blood, The Journal of the American Society of Hematology, vol. 113, no. 11, pp. 2375–2385, 2009.

[19] Y. Shima, M. Yumoto, T. Katsumoto, and I. Kitabayashi, “Mll is essential for nup98-hoxa9-induced leukemia,” Leukemia, vol. 31, no. 10, pp. 2200–2210, 2017.

[20] X. Zhong, A. Prinz, J. Steger, M.-P. Garcia-Cuellar, M. Radsak, A. Bentaher, and R. K. Slany, “Hoxa9 transforms murine myeloid cells by a feedback loop driving expression of key oncogenes and cell cycle control genes,” Blood advances, vol. 2, no. 22, pp. 3137–3148, 2018.

[21] J. Ameijeiras-Alonso, R. M. Crujeiras, and A. Rodrguez-Casal, “multimode: An r package for mode assessment,” árXiv preprint arXiv:1803.00472, 2018.

[22] T. Therneau and T. Lumley, “R survival package,” 2013.

[23] E. Chen, R. K. Schneider, L. J. Breyfogle, E. A. Rosen, L. Poveromo, S. Elf, A. Ko, K. Brumme, R. Levine, B. L. Ebert et al., “Distinct effects of concomitant jak2v617f expression and tet2 loss in mice promote disease progression in myeloproliferative neoplasms,” Blood, The Journal of the American Society of Hematology, vol. 125, no. 2, pp. 327–335, 2015.

[24] Z. Gu, R. Eils, and M. Schlesner, “Complex heatmaps reveal patterns and correlations in multidimensional genomic data,” Bioinformatics, vol. 32, no. 18, pp. 2847–2849, 2016.

[25] T. Chen and C. Guestrin, “Xgboost: A scalable tree boosting system,” in Proceedings of the 22nd acm sigkdd international conference on knowledge discovery and data mining, 2016, pp. 785–794.

[26] D. Benque, S. Bourton, C. Cockerton, B. Cook, J. Fisher, S. Ishtiaq, N. Piterman, A. Taylor, and M. Y. Vardi, “Bma: Visual tool for modeling and analyzing biological networks,” in International Conference on Computer Aided Verification. Springer, 2012, pp. 686–692.

[27] C. G. A. R. Network, “Genomic and epigenomic landscapes of adult de novo acute myeloid leukemia,” New England Journal of Medicine, vol. 368, no. 22, pp. 2059–2074, 2013.

[28] K. Rejlova, A. Musilova, K. S. Kramarzova, M. Zaliova, K. Fiser, M. Alberich-Jorda, J. Trka, and J. Starkova, “Low hox gene expression in pml-rarα-positive leukemia results from suppressed histone demethylation,” Epigenetics, vol. 13, no. 1, pp. 73–84, 2018.

[29] A. Lasa, M. Carnicer, A. Aventin, C. Estivill, S. Brunet, J. Sierra, and J. Nomdedeu, “Meis 1 expression is downregulated through promoter hypermethylation in aml1-eto acute myeloid leukemias,” Leukemia, vol. 18, no. 7, pp. 1231–1237, 2004.

[30] J. C. Byrd, K. Mrózek, R. K. Dodge, A. J. Carroll, C. G. Edwards, D. C. Arthur, M. J. Pettenati, S. R. Patil, K. W. Rao, M. S. Watson et al., “Pretreatment cytogenetic abnormalities are predictive of induction success, cumulative incidence of relapse, and overall survival in adult patients with de novo acute myeloid leukemia: results from cancer and leukemia group b (calgb 8461) presented in part at the 43rd annual meeting of the american society of hematology, orlando, fl, december 10, 2001, and published in abstract form. 59,” Blood, The Journal of the American Society of Hematology, vol. 100, no. 13, pp. 4325–4336, 2002.

[31] Z.-Y. Wang and Z. Chen, “Acute promyelocytic leukemia: from highly fatal to highly curable,” Blood, The Journal of the American Society of Hematology, vol. 111, no. 5, pp. 2505–2515, 2008.

[32] R. L. Levine, M. Wadleigh, J. Cools, B. L. Ebert, G. Wernig, B. J. Huntly, T. J. Boggon, I. Wlodarska, J. J. Clark, S. Moore et al., “Activating mutation in the tyrosine kinase jak2 in polycythemia vera, essential thrombocythemia, and myeloid metaplasia with myelofibrosis,” Cancer cell, vol. 7, no. 4, pp. 387–397, 2005.

[33] F. Delhommeau, S. Dupont, V. D. Valle, C. James, S. Trannoy, A. Masse, O. Kosmider, J.-P. Le Couedic, F. Robert, A. Alberdi et al., “Mutation in tet2 in myeloid cancers,” New England Journal of Medicine, vol. 360, no. 22, pp. 2289–2301, 2009.

[34] S. Chiba, “Dysregulation of tet2 in hematologic malignancies,” International journal of hematology, vol. 105, no. 1, pp. 17–22, 2017.

[35] Y. Z. Paterson, D. Shorthouse, M. W. Pleijzier, N. Piterman, C. Bendtsen, B. A. Hall, and J. Fisher, “A toolbox for discrete modelling of cell signalling dynamics,” Integrative Biology, vol. 10, no. 6, pp. 370–382, 2018.

[36] W. Xiong and J. E. Ferrell, “A positive-feedback-based bistable memory module that governs a cell fate decision,” Nature, vol. 426, no. 6965, pp. 460–465, 2003.

[37] S. Zhao, K. Zoller, M. Masuko, P. Rojnuckarin, X. O. Yang, E. Parganas, K. Kaushansky, J. N. Ihle, T. Papayannopoulou, D. M. Willerford et al., “Jak2, complemented by a second signal from c-kit or flt-3, triggers extensive self-renewal of primary multipotential hemopoietic cells,” The EMBO journal, vol. 21, no. 9, pp. 2159–2167, 2002.

[38] C. E. de Bock, S. Demeyer, S. Degryse, D. Verbeke, B. Sweron, O. Gielen, R. Vandepoel, C. Vicente, M. V. Bempt, A. Dagklis et al., “Hoxa9 cooperates with activated jak/stat signaling to drive leukemia development,” Cancer discovery, vol. 8, no. 5, pp. 616–631, 2018.

[39] L. Bei, C. Shah, H. Wang, W. Huang, L. C. Platanias, and E. A. Eklund, “Regulation of cdx4 gene transcription by hoxa9, hoxa10, the mll-ell oncogene and shp2 during leukemogenesis,” Oncogenesis, vol. 3, no. 12, pp. e135–e135, 2014.

[40] M. T. Bocker, F. Tuorto, G. Raddatz, T. Musch, F.-C. Yang, M. Xu, F. Lyko, and A. Breiling, “Hydroxylation of 5-methylcytosine by tet2 maintains the active state of the mammalian hoxa cluster,” Nature communications, vol. 3, no. 1, pp. 1–12, 2012.

[41] A. Klinakis, C. Lobry, O. Abdel-Wahab, P. Oh, H. Haeno, S. Buonamici, I. van De Walle, S. Cathelin, T. Trimarchi, E. Araldi et al., “A novel tumour-suppressor function for the notch pathway in myeloid leukaemia,” Nature, vol. 473, no. 7346, pp. 230–233, 2011.

[42] P. Chastagner, A. Israel, and C. Brou, “Aip4/itch regulates notch receptor degradation in the absence of ligand,” PloS one, vol. 3, no. 7, 2008.

[43] E. Gallagher, M. Gao, Y.-C. Liu, and M. Karin, “Activation of the e3 ubiquitin ligase itch through a phosphorylation-induced conformational change,” Proceedings of the National Academy of Sciences, vol. 103, no. 6, pp. 1717–1722, 2006.

[44] R. M. de Freitas and C. M. da Costa Maranduba, “Myeloproliferative neoplasms and the jak/stat signaling pathway: an overview,” Revista brasileira de hematologia e hemoterapia, vol. 37, no. 5, pp. 348–353, 2015.

[45] T. Bellon, D. Perrotti, and B. Calabretta, “Granulocytic differentiation of normal hematopoietic precursor cells induced by transcription factor pu. 1 correlates with negative regulation of the c-myb promoter,” Blood, The Journal of the American Society of Hematology, vol. 90, no. 5, pp. 1828–1839, 1997.

[46] W. Zhao, C. Kitidis, M. D. Fleming, H. F. Lodish, and S. Ghaffari, “Erythropoietin stimulates phosphorylation and activation of gata-1 via the pi3-kinase/akt signaling pathway,” Blood, vol. 107, no. 3, pp. 907–915, 2006.

[47] M. Ichikawa, T. Asai, T. Saito, G. Yamamoto, S. Seo, I. Yamazaki, T. Yamagata, K. Mitani, S. Chiba, H. Hirai et al., “Aml-1 is required for megakaryocytic maturation and lymphocytic differentiation, but not for maintenance of hematopoietic stem cells in adult hematopoiesis,” Nature medicine, vol. 10, no. 3, pp. 299–304, 2004.

[48] M. Ichikawa, S. Goyama, T. Asai, M. Kawazu, M. Nakagawa, M. Takeshita, S. Chiba, S. Ogawa, and M. Kurokawa, “Aml1/runx1 negatively regulates quiescent hematopoietic stem cells in adult hematopoiesis,” The Journal of Immunology, vol. 180, no. 7, pp. 4402–4408, 2008.

[49] J. J. Jeong, X. Gu, J. Nie, S. Sundaravel, H. Liu, W.-L. Kuo, T. D. Bhagat, K. Pradhan, J. Cao, S. Nischal et al., “Cytokine-regulated phosphorylation and activation of tet2 by jak2 in hematopoiesis,” Cancer discovery, vol. 9, no. 6, pp. 778–795, 2019.

[50] F. Lohmann and J. J. Bieker, “Activation of eklf expression during hematopoiesis by gata2 and smad5 prior to erythroid commitment,” Development, vol. 135, no. 12, pp. 2071–2082, 2008.

[51] F. Cervantes, F. Passamonti, and G. Barosi, “Life expectancy and prognostic factors in the classic bcr/abl-negative myeloproliferative disorders,” Leukemia, vol. 22, no. 5, pp. 905–914, 2008.

[52] J. C. Aster, W. S. Pear, and S. C. Blacklow, “The varied roles of notch in cancer,” Annual Review of Pathology: Mechanisms of Disease, vol. 12, pp. 245–275, 2017.

[53] F. P. Lampreia, J. G. Carmelo, and F. Anjos-Afonso, “Notch signaling in the regulation of hematopoietic stem cell,” Current stem cell reports, vol. 3, no. 3, pp. 202–209, 2017.

