## Supplementary Text for "*HOXA9* acts as a regulatory switch in acute myeloid leukaemia and myeloproliferative neoplasms"

Talarmain et al.

[*RUNX1*, MYB and MYC general expression trend in [36] are consistent with our model.	6](#_Toc64978796)

### Supplemental Methods

#### Qualitative Networks

Qualitative networks are an extension of Boolean networks described in detail in [1]. Of note, the nodes representing molecular expression are discrete variables with more than two states. This means higher resolution in expression levels. Additionally, the activity of a variable is determined by an algebraic target function rather than a table of values. The formalism is defined below.

A Qualitative network $Q(C,T,N)$ consists of components in $C$ which can take values in $\{0,1,..,N\}.$ $N$ is a constant integer with any possible values above one, and is called the node granularity. If $N$ is one, the model is a Boolean network. $T$ is the set of target functions, which are the functions that determines the level toward which each component moves at the following time step. At each time step, each component moves by a maximum of one level. The update of each component $c_{i}\in C$ can therefore be mathematically defined as:

$$c_{i}(t+1) = \left\{ \begin{aligned} c_{i} + 1\text{ if} {target}_{i}(s(t)) > c_{i} \\ c_{i} - 1 \text{if} {target}_{i}(s(t)) < c_{i} \\ c_{i} \text{if} {target}_{i}(s(t)) = c_{i} \end{aligned} \right.$$

with $s(t)$ the current state of the network and ${target}_{i}\in T$ the target function of $c_{i}$. ${target}_{i}$returns a value in $\{0,1,..,N\}$. The default target function is the difference between the amount of activation ${act}_{i}$ and the amount of inhibition ${inh}_{i}$ a component $c_{i}$ is subject to. Both are averaged by the total number of activating/inhibiting components and are defined as follows:

$${act}_{i} = \frac{\sum_{\alpha_{ji}>0} \alpha_{ji}c_{j}}{\sum_{\alpha_{ji}>0} \alpha_{ji}}$$

$${inh}_{i} = \frac{\sum_{\alpha_{ji}<0} \alpha_{ji}c_{j}}{\sum_{\alpha_{ji}<0} \alpha_{ji}}$$

All models are built using the BioModelAnalyzer (BMA) tool [2] (https://biomodelanalyzer.org/), and analysed using LTL [3] and stability analysis [4]. BMA includes two types of edges representing respectively the activation or the inhibition of a node.

#### Computational network model construction: description and literature review

To build and validate a computational network model, the process must iterate over three fundamental steps:

- Develop a “specification”- a set of specific biological phenotypes and gene activities associated with each input condition (for example, the genotype or environment)
- Create a network using genes of interest and their interactions as nodes and edges. Phenotypes are also modelled as nodes in the network
- Refine the “target functions” on each gene and phenotype that govern how the gene responds to the activity of upstream element

A “correct” model will match all input specifications. As the model is expanded to include extended specifications or more genes, this cycle of testing and refinement must be repeated.

The MPN model includes four genotypes in the specifications; wild-type JAK2/TET2, TET2 loss of function, JAK2 gain of function, and the JAK2/TET2 double mutant (Table 1). Variables are discrete and range from zero to two, representing inactive, resting, and hyperactive states.

#### Gene network construction

Genes included in the network were selected based on their relationships with the three core genes (TET2, JAK2, and HOXA9), and their involvement in control of modelled phenotypes. *SPI1* and *CEBPA* have well defined roles in monocyte and macrophage lineages [5]. *SPI1* activates *CEBPA* in early progenitors [6, 7] and *CEBPA* helps the transition from CMP to GMP [8]. Both genes are essential for myeloid differentiation and are downregulators of progenitor proliferation.

Differentiation is associated with a reduced cell proliferation [9]. Our model includes *MYB* and *E2F1* as upstream effectors for CMP and GMP expansion [10, 11] which are both inactivated by *SPI1* and *CEBPA* [12-14]. Furthermore, we include *GATA1* in our network as part of JAK2 pathway and its role in determining the erythroid lineage. Both GATA1 and JAK2 genes have been shown to be important players in MEP progenitor production and erythropoiesis [15, 16]. *GATA1* is additionally required for *KLF1* activation, a marker of erythroid differentiation [17]. KLF1 has also been established as a downstream target of phosphorylated *TET2* in erythroid cell lines and *TET2* phosphorylation is induced by *JAK2* [18]. *JAK2* is an upstream regulator of *GATA1* via AKT [19], but also plays an role in MEP expansion with *STAT5* and MAPK activation of the anti-apoptotic gene *BCL2L1* [20, 21].

Whilst *JAK2* is associated with erythroid differentiation, it also can play a role in determining the myeloid lineage through *STAT3* mediated activation of *SPI1* [22].

Finally, *SPI1* and *GATA1* mutually inhibit one another [23]. This feature is necessary for erythroid/myeloid lineage commitment [24]. We additionally include *RUNX1* as a link between those hematopoietic genes and our *JAK2*/*TET2*/HOXA9 motif through *RUNX1* activation by HOXA9. *RUNX1* upregulation has been associated with *HOXA9* upregulation in early stem and progenitor cells [25, 26] and therefore is a strong candidate to link our motif with our hematopoietic genetic regulators. *RUNX1* is found in the earliest stages of haematopoiesis which makes this gene essential for a fully functional haematopoiesis [27]. We therefore link this gene to the rest of our network through its downstream targets *SPI1* for the myeloid lineage [28] and with *GATA1* for the erythroid lineage [29, 30].

#### Model genotype/phenotype specifications

The model specifications are defined in Table 1 in the paper and are described in more details in this section. In the wild-type state, the model is stable with all variables (genes and phenotypes) equal to one. In the single *TET2* loss-of-function mutation state, the cell has increased stem cell self-renewal [31], elevated CMP expansion [32] and diminished overall differentiation [33] with a skew towards the granulocyte-monocyte lineage [34]. This is tested in our model by looking for stability, with an increase in GMP expansion, whilst MEP expansion remains at its wild type levels. In the JAK2 mutant (overactivation) background, erythroid differentiation and MEP expansion are both increased [35, 36]. Furthermore, GMP expansion is also increased to reflect *JAK2* stimulation of myeloid cells, but GMP differentiation remains at wild-type levels as the erythroid lineage specifically is preferred [37].

Finally, the double mutant is a bifurcating system with two fixed points and no cyclic attractors. Each fixed point represents alternative orders of TET2/JAK2 mutations; that is to say, *TET2* first or *JAK2* first double mutants. We characterised those states using Ortmann et al [38]. Both fixed points have an increased stem cell self-renewal due to *TET2* loss. GMP and MEP expansion are also increased in both states as *TET2* loss promotes granulocyte-monocyte development, while *JAK2* favours the erythroid lineage. However, following observations in Ortmann et al [38], in our model *TET2* first mutants have an increased CMP expansion not observed in *JAK2* first. Additionally, the differentiation loss in *JAK2* single mutants is retrieved by *JAK2* overexpression, resulting in GMP differentiation being in its normal state in both double mutants. Experiments have shown that *JAK2* first double mutant patients have an increased number of mature erythroid cells, and so erythroid differentiation is increased in *JAK2* first but not *TET2* first double mutant. In total, both double mutants share four common phenotypes. Only CMP expansion and erythroid differentiation differ between the two fixed points consistent with patient characteristics described in Ortmann et al [38].

#### *RUNX1*, MYB and MYC general expression trend in [36] are consistent with our model.

We plot *RUNX1, MYB* and *MYC* expression for the different genotypes in the data and compare them to their expression in our biological network. In Figure S1, S2 and S3, all genotypes come from [36] and therefore the genotypes "JAK2" and "TET2" refer to the single mutant mouse models, and "DM" is the double mutant with *JAK2* mutated first. "WT" designates the wild type (no mutation) genotype.

#### XGBoost

We use XGBoost (e**X**treme **G**radient **Boost**ing) to rank different gene pathways that have been well described in cancer to identify which pathways and genes amongst these pathways have the highest correlation with a gene of interest and its expression level in the AML patients (TCGA RNASeq) [39]. We start by splitting the patients into two groups: one with the highest expression for our gene of interest and another with the lowest expression. We use XGBoost binary classification to determine which pathways are the best to classify patients into the right cohort. Fifteen text files containing different subset of genes corresponding to popular cancer pathways were previously generated using literature: cell cycle, EMT, HIPPO, MAPK, MYC, NFKB, NOTCH, NRF2, PI3K, RAS-RTK, TGF-β, TP53, WNT, WNT downstream and WNT drivers. We change the default evaluation metrics in XGBoost to the logarithmic loss (logloss) function. We also tune the *colsample_bytree* parameter to 0.3 instead of 1 (default value). By setting this value to 0.3, we ask XGBoost to build trees with randomly picked 30% genes per pathway. Therefore, it excludes in some iterations the genes that could be highly correlated to our gene of interest and therefore reduce the accuracy score if other genes of the pathway are not good classifiers. Finally, we use a Matthew Correlation Coefficient (*MCC*) to score the competence of each pathway to classify our patients into the right cohort:

$$MCC= \frac{TP \times TN-FP \times FN}{TP+TN+FP+FN}$$

True Positive ($TP$) and True Negative ($TN$) are counts representing how many times a pathway has correctly classified a patient into respectively the high or low cohort. False Positive ($FP$) represents the number of times a pathway has classified a patient with low expression into the high cohort, and vice versa for the False Negative ($FN$). $MCC$ score varies -1 and 1, but we translate it into percentage in our figures.

We use SHAP to explain the output of our XGBoost models. SHAP (**SH**apley **A**dditive ex**P**lanations) aims to ease the interpretability of complex models by representing the importance of model features with shapley values [40]. Documentation on SHAP and how to use it can be found on github (http://github.com/slundberg/shap).

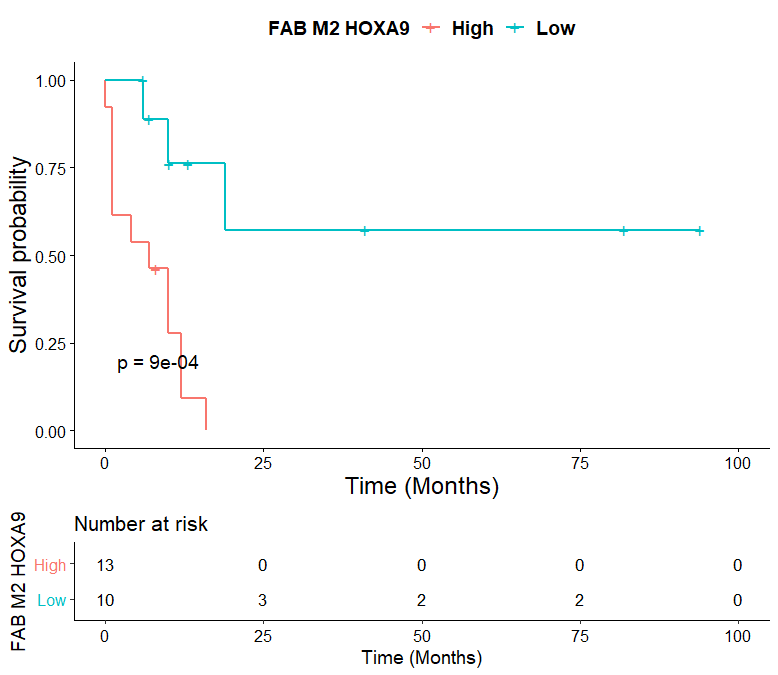

### Supplemental Figures

**Figure S1. *HOXA9* high and low cohorts of M2 AML patients show distinct survival probabilities.** None of the 13 patients with high expression for *HOXA9* survive past 20 months while three among the 10 patients with low expression reach 25 months.

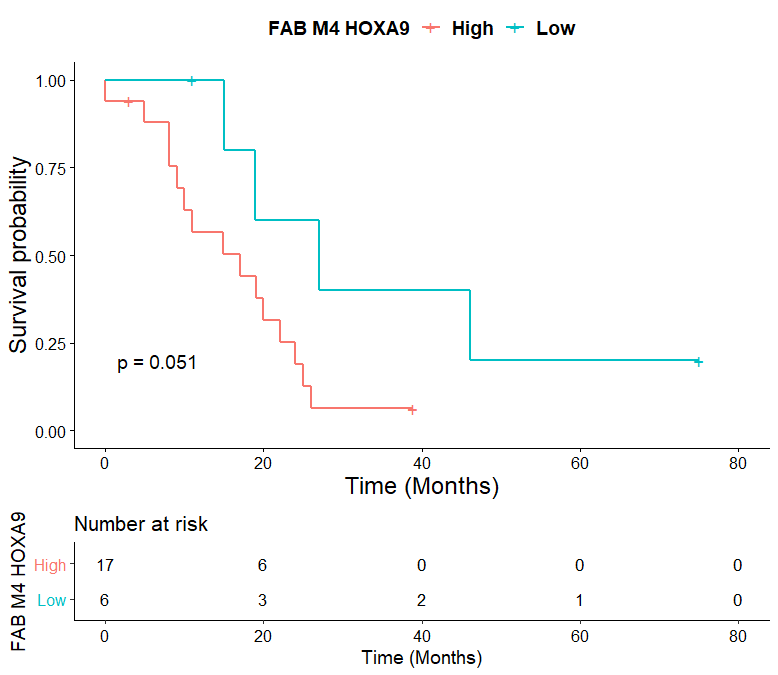
**Figure S2. *HOXA9* high and low cohorts of M4 AML patients show distinct survival probabilities.** The Kaplan Meier curves for the survival of M4 patients indicates a trend towards lower survival probability for patients with high *HOXA9* expression compared to the low-*HOXA9* cohort ($p=0.051$).

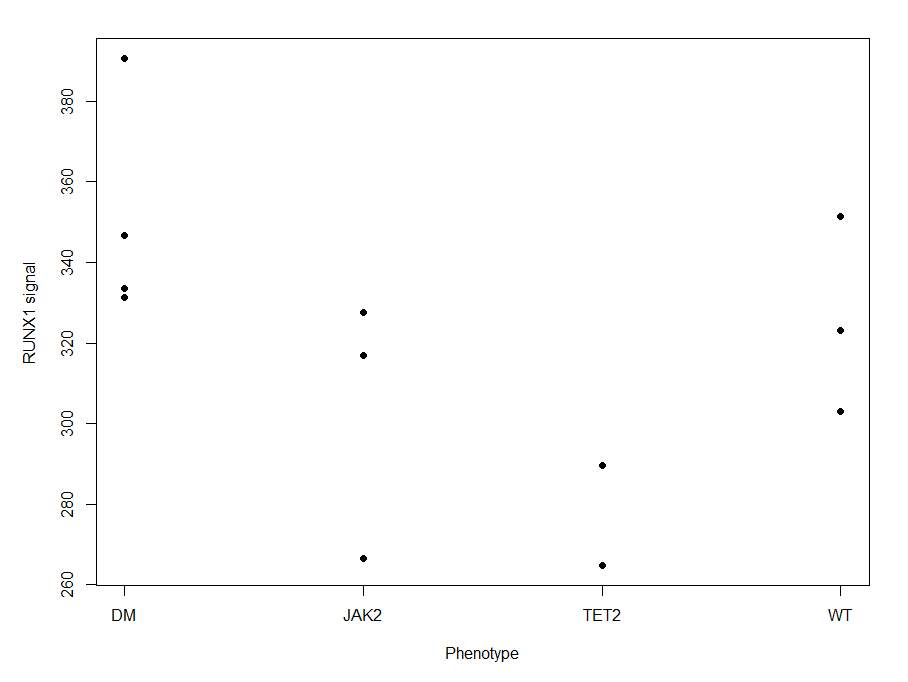

**Figure S3. *RUNX1* expression in public MPN data share the same trend than our model.** Our model expects *RUNX1* to be higher in the *JAK2* single and *JAK2* first double mutants and lower in *TET2* single mutants compared to the wild type. Despite the low number of data points, the trends for *RUNX1* expression in the different genotypes are consistent with our findings.

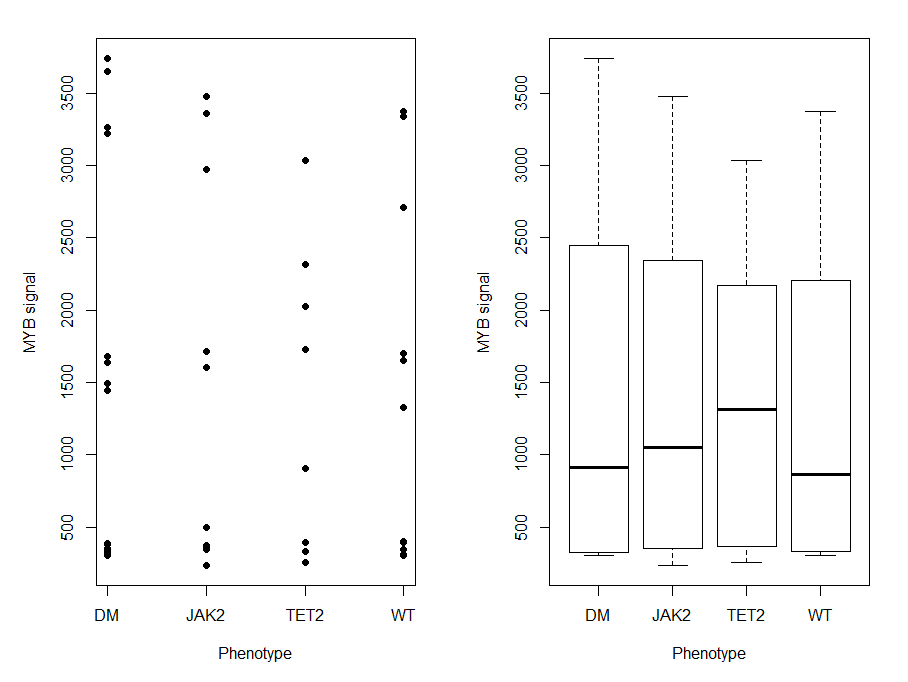

**Figure S4.** ***MYB* expression in public MPN data share the same trend than our model.** Our model expects *MYB* expression to be higher in the *TET2* single mutants while *JAK2* single and *JAK2* first double mutants have a similar *MYB* expression compared to the wild type genotype. Despite the low number of data points, using the boxplot figure we identify that the trends for *MYB* expression in the different genotypes are consistent with our findings.

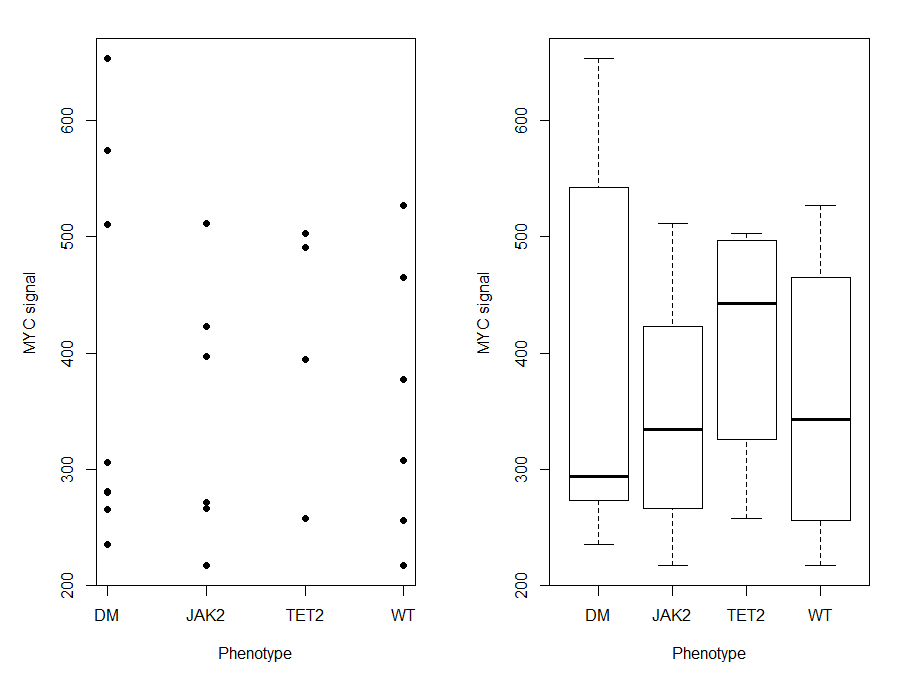

**Figure S5. *MYC* expression in public MPN data share the same trend than our model.** Our model expects *MYC* expression to be higher in the *TET2* single mutants while *JAK2* single and *JAK2* first double mutants should show a lower expression compared the wild type genotype. Despite the low number of data points, using the boxplot figure we identify that the trends for *MYC* expression in the different genotypes fit our findings for at least the *TET2* single mutant and the *JAK2* first double mutant.

### Supplemental Tables

| **Upstream gene** | **Interaction type** | **Downstream gene** | **Reference** |
| --- | --- | --- | --- |
| *TET2* | Activates | *HOXA9* | [41] |
| *JAK2* | Activates | *HOXA9* | [42] |
| *HOXA9* | Activates | *HOXA9* | [43] |
| *HOXA9* | Activates | *RUNX1* | [25, 26] |
| *RUNX1* | Activates | *SPI1* | [28] |
| *RUNX1* | Activates | *GATA1* | [29, 30] |
| *SPI1* | Inhibits | *GATA1* | [23] |
| *GATA1* | Inhibits | *SPI1* | [23] |
| *SPI1* | Activates | *CEBPa* | [6] |
| *CEBPa* | Inhibits | *E2F1* | [13] |
| *CEBPa* | Activates | GMP differentiation | [8] |
| *E2F1* | Activates | GMP expansion | [11] |
| *RUNX1* | Inhibits | *MYB* | Model prediction |
| *MYB* | Activates | CMP expansion | [10] |
| *TET2* | Activates | SC self renewal | [31] |
| *GATA1* | Activates | *KLF1* | [17] |
| *GATA1* | Inhibits | *MYC* | [44] |
| *MYC* | Activates | MEP expansion | [44] |
| *KLF1* | Activates | Erythroid differentiation | [17] |
| *TET2* | Activates | *TET2p* | [18] |
| *JAK2* | Activates | *TET2p* | [18] |
| *TET2p* | Activates | *KLF1* | [18] |
| *JAK2* | Activates | *STAT3* | [45] |
| *JAK2* | Activates | *STAT5* | [45] |
| *STAT3* | Activates | *SPI1* | [22] |
| *STAT5* | Activates | *AKT* | [19] |
| *AKT* | Activates | *GATA1* | [19] |
| *STAT5* | Activates | *MAPK* | [20] |
| *MAPK* | Activates | *BCL2L1* | [20] |
| *BCL2L1* | Activates | MEP expansion | [21] |
| *TET2* | Activates | MEP expansion | [34] |
| *MAPK* | Activates | *JNK1* | [46] |
| *JNK1* | Activates | *ITCH* | [47] |
| *ITCH* | Inhibits | *NOTCH* | [48] |
| *NOTCH* | Inhibits | GMP expansion | [49] |

**Table S1. Gene interaction table for *JAK2*/*TET2* BMA model.**

| **Node** | **Target Function** | **References** | **Comments** |
| --- | --- | --- | --- |
| HOXA9 | JAK2*TET2 + 2*max(0,(HOXA9-1))*max(0,JAK2-1) | [42],[41],[43] | Activation by *JAK2*, *TET2* and itself. Memory property prediction. |
| SPI1 | STAT3 + min( (RUNX1-1) , (1-GATA1) ) | [28], [23],[19] | Activation by *STAT3* and *RUNX1* and inhibition loop with *GATA1*. Minimum function to have normal *SPI1* expression when *GATA1* is underexpressed but decreased expression when *GATA1* is overexpressed independently of *RUNX1*. |
| GATA1 | AKT+RUNX1-max(SPI1,1) | [29, 23],[22] | Activation by *AKT* and *RUNX1* and inhibition loop with *SPI1*. Maximum function necessary as loss of *SPI1* does not increase erythroid differentiation. |
| MEP expansion | min( BCL2L1+TET2 , max( MYC, BCL2L1) ) | [36],[44],[21],[34],[35] | *TET2* loss reduces erythroid progenitors by skewing toward myeloid lineage. *JAK2* positive regulation of the erythroid lineage is stronger than *TET2* through *MYC* activation. |

**Table S2. Target functions of the model variables.** Except for the nodes indicated by this table, all nodes have the default target function of the BioModelAnalyzer (BMA) tool which is the difference between the average state of all the variables activating the current node and the average of all the variables inhibiting it. If there is no activating variable in the model, the target function equals the difference between a constant representing the basal activity of the node and the inhibiting variables. The constant is calculated so that in the healthy state (no mutation), all variables equal 1.

### Supplemental References

[1]	M. A. Schaub, T. A. Henzinger, and J. Fisher, “Qualitative networks: a symbolic approach to analyze biological signaling networks,” *BMC systems biology*, vol. 1, no. 1, p. 4, 2007.

[2]	D. Benque, S. Bourton, C. Cockerton, B. Cook, J. Fisher, S. Ishtiaq, N. Piterman, A. Taylor, and M. Y. Vardi, “Bma: Visual tool for modeling and analyzing biological networks,” in *International Conference on Computer Aided Verification*. Springer, 2012, pp. 686–692.

[3]	A. Pnueli, “The temporal logic of programs,” in *18th Annual Symposium on Foundations of Computer Science (sfcs 1977)*. IEEE, 1977, pp. 46–57.

[4]	B. Cook, J. Fisher, E. Krepska, and N. Piterman, “Proving stabilization of biological systems,” in *International Workshop on Verification, Model Checking, and Abstract Interpretation*. Springer, 2011, pp. 134–149.

[5]	F. Rosenbauer and D. G. Tenen, “Transcription factors in myeloid development: balancing differentiation with transformation,” *Nature Reviews Immunology*, vol. 7, no. 2, pp. 105–117, 2007.

[6]	P. Burda, N. Curik, J. Kokavec, P. Basova, D. Mikulenkova, A. I. Skoultchi, J. Zavadil, and T. Stopka, “Pu. 1 activation relieves gata-1–mediated repression of cebpa and cbfb during leukemia differentiation,” *Molecular Cancer Research*, vol. 7, no. 10, pp. 1693–1703, 2009.

[7]	S. Pundhir, F. K. B. Lauridsen, M. B. Schuster, J. S. Jakobsen, Y. Ge, E. M. Schoof, N. Rapin, J. Waage, M. S. Hasemann, and B. T. Porse, “Enhancer and transcription factor dynamics during myeloid differentiation reveal an early differentiation block in cebpa null progenitors,” *Cell reports*, vol. 23, no. 9, pp. 2744–2757, 2018.

[8]	P. Zhang, J. Iwasaki-Arai, H. Iwasaki, M. L. Fenyus, T. Dayaram, B. M. Owens, H. Shigematsu, E. Levantini, C. S. Huettner, J. A. Lekstrom-Himes *et al.*, “Enhancement of hematopoietic stem cell repopulating capacity and self-renewal in the absence of the transcription factor c/ebpα,” *Immunity*, vol. 21, no. 6, pp. 853–863, 2004.

[9]	A. D. Friedman, “Runx1, c-myb, and c/ebpα couple differentiation to proliferation or growth arrest during hematopoiesis,” *Journal of cellular biochemistry*, vol. 86, no. 4, pp. 624–629, 2002.

[10]	Y. K. Lieu and E. P. Reddy, “Impaired adult myeloid progenitor cmp and gmp cell function in conditional c-myb-knockout mice,” *Cell Cycle*, vol. 11, no. 18, pp. 3504–3512, 2012.

[11]	B. T. Porse, D. Bryder, K. Theilgaard-Monch, M. S. Hasemann, K. Anderson, I. Damgaard, S. E. W. Jacobsen, and C. Nerlov, “Loss of c/ebpα cell cycle control increases myeloid progenitor proliferation and transforms the neutrophil granulocyte lineage,” *The Journal of experimental medicine*, vol. 202, no. 1, pp. 85–96, 2005.

[12]	T. Bellon, D. Perrotti, and B. Calabretta, “Granulocytic differentiation of normal hematopoietic precursor cells induced by transcription factor pu. 1 correlates with negative regulation of the c-myb promoter,” *Blood, The Journal of the American Society of Hematology*, vol. 90, no. 5, pp. 1828–1839, 1997.

[13]	B. T. Porse, T. Å. Pedersen, X. Xu, B. Lindberg, U. M. Wewer, L. Friis-Hansen, and C. Nerlov, “E2f repression by c/ebpα is required for adipogenesis and granulopoiesis in vivo,” *Cell*, vol. 107, no. 2, pp. 247–258, 2001.

[14]	L. M. Johansen, A. Iwama, T. A. Lodie, K. Sasaki, D. W. Felsher, T. R. Golub, and D. G. Tenen, “c-myc is a critical target for c/ebpα in granulopoiesis,” *Molecular and cellular biology*, vol. 21, no. 11, pp. 3789–3806, 2001.

[15]	X. Han, J. Zhang, Y. Peng, M. Peng, X. Chen, H. Chen, J. Song, X. Hu, M. Ye, J. Li *et al.*, “Unexpected role for p19ink4d in posttranscriptional regulation of gata1 and modulation of human terminal erythropoiesis,” *Blood, The Journal of the American Society of Hematology*, vol. 129, no. 2, pp. 226–237, 2017.

[16]	H. Neubauer, A. Cumano, M. Müller, H. Wu, U. Huffstadt, and K. Pfeffer, “Jak2 deficiency defines an essentialdevelopmental checkpoint in definitivehematopoiesis,” *Cell*, vol. 93, no. 3, pp. 397–409, 1998.

[17]	F. Lohmann and J. J. Bieker, “Activation of eklf expression during hematopoiesis by gata2 and smad5 prior to erythroid commitment,” *Development*, vol. 135, no. 12, pp. 2071–2082, 2008.

[18]	J. J. Jeong, X. Gu, J. Nie, S. Sundaravel, H. Liu, W.-L. Kuo, T. D. Bhagat, K. Pradhan, J. Cao, S. Nischal *et al.*, “Cytokine-regulated phosphorylation and activation of tet2 by jak2 in hematopoiesis,” *Cancer discovery*, vol. 9, no. 6, pp. 778–795, 2019.

[19]	I. Geron, A. E. Abrahamsson, C. F. Barroga, E. Kavalerchik, J. Gotlib, J. D. Hood, J. Durocher, C. C. Mak, G. Noronha, R. M. Soll *et al.*, “Selective inhibition of jak2-driven erythroid differentiation of polycythemia vera progenitors,” *Cancer cell*, vol. 13, no. 4, pp. 321–330, 2008.

[20]	M. Socolovsky, A. E. Fallon, C. Brugnara, and H. F. Lodish, “Fetal anemia and apoptosis of red cell progenitors in stat5a-/- 5b-/- mice: a direct role for stat5 in bcl-xl induction,” *Cell*, vol. 98, no. 2, pp. 181–191, 1999.

[21]	M. Mori, M. Uchida, T. Watanabe, K. Kirito, K. Hatake, K. Ozawa, and N. Komatsu, “Activation of extracellular signal-regulated kinases erk1 and erk2 induces bcl-xl up-regulation via inhibition of caspase activities in erythropoietin signaling,” *Journal of cellular physiology*, vol. 195, no. 2, pp. 290–297, 2003.

[22]	A. D. Panopoulos, D. Bartos, L. Zhang, and S. S. Watowich, “Control of myeloid-specific integrin αmβ2 (cd11b/cd18) expression by cytokines is regulated by stat3-dependent activation of pu. 1,” *Journal of Biological Chemistry*, vol. 277, no. 21, pp. 19001–19007, 2002.

[23]	P. Zhang, G. Behre, J. Pan, A. Iwama, N. Wara-Aswapati, H. S. Radomska, P. E. Auron, D. G. Tenen, and Z. Sun, “Negative cross-talk between hematopoietic regulators: Gata proteins repress pu. 1,” *Proceedings of the National Academy of Sciences*, vol. 96, no. 15, pp. 8705–8710, 1999.

[24]	P. Burda, P. Laslo, and T. Stopka, “The role of pu. 1 and gata-1 transcription factors during normal and leukemogenic hematopoiesis,” *Leukemia*, vol. 24, no. 7, pp. 1249–1257, 2010.

[25]	S. Tsuzuki and M. Seto, “Expansion of functionally defined mouse hematopoietic stem and progenitor cells by a short isoform of runx1/aml1,” *Blood, The Journal of the American Society of Hematology*, vol. 119, no. 3, pp. 727–735, 2012.

[26]	V. Azcoitia, M. Aracil, C. Martnez-A, and M. Torres, “The homeodomain protein meis1 is essential for definitive hematopoiesis and vascular patterning in the mouse embryo,” *Developmental biology*, vol. 280, no. 2, pp. 307–320, 2005.

[27]	J. D. Growney, H. Shigematsu, Z. Li, B. H. Lee, J. Adelsperger, R. Rowan, D. P. Curley, J. L. Kutok, K. Akashi, I. R. Williams *et al.*, “Loss of runx1 perturbs adult hematopoiesis and is associated with a myeloproliferative phenotype,” *Blood*, vol. 106, no. 2, pp. 494–504, 2005.

[28]	Y. Huang, K. Sitwala, J. Bronstein, D. Sanders, M. Dandekar, C. Collins, G. Robertson, J. MacDonald, T. Cezard, M. Bilenky *et al.*, “Identification and characterization of hoxa9 binding sites in hematopoietic cells,” *Blood, The Journal of the American Society of Hematology*, vol. 119, no. 2, pp. 388–398, 2012.

[29]	K. E. Elagib, F. K. Racke, M. Mogass, R. Khetawat, L. L. Delehanty, and A. N. Goldfarb, “Runx1 and gata-1 coexpression and cooperation in megakaryocytic differentiation,” *Blood*, vol. 101, no. 11, pp. 4333–4341, 2003.

[30]	T. Yokomizo, K. Hasegawa, H. Ishitobi, M. Osato, M. Ema, Y. Ito, M. Yamamoto, and S. Takahashi, “Runx1 is involved in primitive erythropoiesis in the mouse,” *Blood, The Journal of the American Society of Hematology*, vol. 111, no. 8, pp. 4075–4080, 2008.

[31]	L. Cimmino, I. Dolgalev, Y. Wang, A. Yoshimi, G. H. Martin, J. Wang, V. Ng, B. Xia, M. T. Witkowski, M. Mitchell-Flack *et al.*, “Restoration of tet2 function blocks aberrant self-renewal and leukemia progression,” *Cell*, vol. 170, no. 6, pp. 1079–1095, 2017.

[32]	H. Kunimoto, Y. Fukuchi, M. Sakurai, K. Sadahira, Y. Ikeda, S. Okamoto, and H. Nakajima, “Tet2 disruption leads to enhanced self-renewal and altered differentiation of fetal liver hematopoietic stem cells,” *Scientific reports*, vol. 2, p. 273, 2012.

[33]	M. Ko, H. S. Bandukwala, J. An, E. D. Lamperti, E. C. Thompson, R. Hastie, A. Tsangaratou, K. Rajewsky, S. B. Koralov, and A. Rao, “Ten-eleven-translocation 2 (tet2) negatively regulates homeostasis and differentiation of hematopoietic stem cells in mice,” *Proceedings of the National Academy of Sciences*, vol. 108, no. 35, pp. 14566–14571, 2011.

[34]	K. Moran-Crusio, L. Reavie, A. Shih, O. Abdel-Wahab, D. Ndiaye-Lobry, C. Lobry, M. E. Figueroa, A. Vasanthakumar, J. Patel, X. Zhao *et al.*, “Tet2 loss leads to increased hematopoietic stem cell self-renewal and myeloid transformation,” *Cancer cell*, vol. 20, no. 1, pp. 11–24, 2011.

[35]	C. H. Jamieson, J. Gotlib, J. A. Durocher, M. P. Chao, M. R. Mariappan, M. Lay, C. Jones, J. L. Zehnder, S. L. Lilleberg, and I. L. Weissman, “The jak2 v617f mutation occurs in hematopoietic stem cells in polycythemia vera and predisposes toward erythroid differentiation,” *Proceedings of the National Academy of Sciences*, vol. 103, no. 16, pp. 6224–6229, 2006.

[36]	E. Chen, R. K. Schneider, L. J. Breyfogle, E. A. Rosen, L. Poveromo, S. Elf, A. Ko, K. Brumme, R. Levine, B. L. Ebert *et al.*, “Distinct effects of concomitant jak2v617f expression and tet2 loss in mice promote disease progression in myeloproliferative neoplasms,” *Blood, The Journal of the American Society of Hematology*, vol. 125, no. 2, pp. 327–335, 2015.

[37]	H. Akada, D. Yan, H. Zou, S. Fiering, R. E. Hutchison, and M. G. Mohi, “Conditional expression of heterozygous or homozygous jak2v617f from its endogenous promoter induces a polycythemia vera–like disease,” *Blood, The Journal of the American Society of Hematology*, vol. 115, no. 17, pp. 3589–3597, 2010.

[38]	C. A. Ortmann, D. G. Kent, J. Nangalia, Y. Silber, D. C. Wedge, J. Grinfeld, E. J. Baxter, C. E. Massie, E. Papaemmanuil, S. Menon *et al.*, “Effect of mutation order on myeloproliferative neoplasms,” *New England Journal of Medicine*, vol. 372, no. 7, pp. 601–612, 2015.

[39]	T. Chen and C. Guestrin, “Xgboost: A scalable tree boosting system,” in *Proceedings of the 22nd acm sigkdd international conference on knowledge discovery and data mining*, 2016, pp. 785–794.

[40]	S. M. Lundberg and S.-I. Lee, “A unified approach to interpreting model predictions,” in *Advances in neural information processing systems*, 2017, pp. 4765–4774.

[41]	M. T. Bocker, F. Tuorto, G. Raddatz, T. Musch, F.-C. Yang, M. Xu, F. Lyko, and A. Breiling, “Hydroxylation of 5-methylcytosine by tet2 maintains the active state of the mammalian hoxa cluster,” *Nature communications*, vol. 3, no. 1, pp. 1–12, 2012.

[42]	L. Bei, C. Shah, H. Wang, W. Huang, L. C. Platanias, and E. A. Eklund, “Regulation of cdx4 gene transcription by hoxa9, hoxa10, the mll-ell oncogene and shp2 during leukemogenesis,” *Oncogenesis*, vol. 3, no. 12, pp. e135–e135, 2014.

[43]	X. Zhong, A. Prinz, J. Steger, M.-P. Garcia-Cuellar, M. Radsak, A. Bentaher, and R. K. Slany, “Hoxa9 transforms murine myeloid cells by a feedback loop driving expression of key oncogenes and cell cycle control genes,” *Blood advances*, vol. 2, no. 22, pp. 3137–3148, 2018.

[44]	V. Munugalavadla, L. C. Dore, B. L. Tan, L. Hong, M. Vishnu, M. J. Weiss, and R. Kapur, “Repression of c-kit and its downstream substrates by gata-1 inhibits cell proliferation during erythroid maturation,” *Molecular and cellular biology*, vol. 25, no. 15, pp. 6747–6759, 2005.

[45]	W. J. Leonard and J. J. O’Shea, “Jaks and stats: biological implications,” *Annual review of immunology*, vol. 16, no. 1, pp. 293–322, 1998.

[46]	H.-L. Huang, M.-J. Hsieh, M.-H. Chien, H.-Y. Chen, S.-F. Yang, and P.-C. Hsiao, “Glabridin mediate caspases activation and induces apoptosis through jnk1/2 and p38 mapk pathway in human promyelocytic leukemia cells,” *PLoS One*, vol. 9, no. 6, p. e98943, 2014.

[47]	E. Gallagher, M. Gao, Y.-C. Liu, and M. Karin, “Activation of the e3 ubiquitin ligase itch through a phosphorylation-induced conformational change,” *Proceedings of the National Academy of Sciences*, vol. 103, no. 6, pp. 1717–1722, 2006.

[48]	P. Chastagner, A. Israel, and C. Brou, “Aip4/itch regulates notch receptor degradation in the absence of ligand,” *PloS one*, vol. 3, no. 7, 2008.

[49]	A. Klinakis, C. Lobry, O. Abdel-Wahab, P. Oh, H. Haeno, S. Buonamici, I. van De Walle, S. Cathelin, T. Trimarchi, E. Araldi *et al.*, “A novel tumour-suppressor function for the notch pathway in myeloid leukaemia,” *Nature*, vol. 473, no. 7346, pp. 230–233, 2011.
